# Direct Comparative Analysis of 10X Genomics Chromium and Smart-seq2

**DOI:** 10.1101/615013

**Authors:** Xiliang Wang, Yao He, Qiming Zhang, Xianwen Ren, Zemin Zhang

## Abstract

Single cell RNA sequencing (scRNA-seq) is widely used for profiling transcriptomes of individual cells. The droplet-based 10X Genomics Chromium (10X) approach and the plate-based Smart-seq2 full-length method are two frequently-used scRNA-seq platforms, yet there are only a few thorough and systematic comparisons of their advantages and limitations. Here, by directly comparing the scRNA-seq data by the two platforms from the same samples of CD45-cells, we systematically evaluated their features using a wide spectrum of analysis. Smart-seq2 detected more genes in a cell, especially low abundance transcripts as well as alternatively spliced transcripts, but captured higher proportion of mitochondrial genes. The composite of Smart-seq2 data also resembled bulk RNA-seq data better. For 10X-based data, we observed higher noise for mRNA in the low expression level. Despite the poly(A) enrichment, approximately 10-30% of all detected transcripts by both platforms were from non-coding genes, with lncRNA accounting for a higher proportion in 10X. 10X-based data displayed more severe dropout problem, especially for genes with lower expression levels. However, 10X-data can better detect rare cell types given its ability to cover a large number of cells. In addition, each platform detected different sets of differentially expressed genes between cell clusters, indicating the complementary nature of these technologies. Our comprehensive benchmark analysis offers the basis for selecting the optimal scRNA-seq strategy based on the objectives of each study.

## Introduction

Following the first single-cell RNA sequencing (scRNA-seq) method developed in 2009 [1], scRNA-seq has dramatically influenced many research fields ranging from cancer biology, stem cell biology to immunology [2–5]. Compared with RNA-seq of bulk tissues with millions of cells, scRNA-seq offers the opportunity to dissect the composition of tissues and the dynamic of transcriptional states, as well as to discover rare cell types. With the improvement of sequencing technologies, scRNA-seq is becoming robust and broadly accessible to perform transcriptome analysis [6].

Two scRNA-seq platforms are frequently used [7, 8]: Smart-seq2 [9] and 10X (10X Genomics Chromium, 10X Genomics, Pleasanton, CA). Smart-seq2 is based on microtiter plates [10, 11], where mRNA is isolated and reverse transcribed to cDNA for high-throughput sequencing for each cell [12]. Reads mapped to a gene are used to quantify its expression in each cell, and TPM (Transcripts Per Kilobase Million) is a common metric of expression normalization [13, 14]. By contrast, 10X is a droplet-based scRNA-seq method, allowing genome-wide expression profiling for thousands of cells at once. The UMI (unique molecular identifier) is used to directly quantify the expression level of each gene [15]. Both TPM (Smart-seq2) and normalized UMI (10X) is analyzed to detect HVGs (highly variable genes), which are often used for either cellular phenotype classification or new subpopulation identification [16].

Although each platform has its own expected advantages and drawbacks based on the design of each method, there are only a few systematic comparisons of Smart-seq2 and 10X [17, 18]. Here, we applied these two technologies to the same set of samples, and directly compared the sensitivity (the probability to detect transcripts present in a single cell), precision (variation of the quantification), and power (subpopulation identification) of these two platforms.

## Results

### Data generation and evaluation

Our data were derived from two cancer patients. For the first patient, diagnosed to have hepatocellular carcinoma (HCC), we collected the liver tumor (LT) and its adjacent non-tumor tissue (NT). For the second patient, diagnosed to have rectal cancer with liver metastasis, we collected both the primary tumor (PT) and the metastasized tumor (MT). For each sample, we used fluorescence activated cell sorting (FACS) to obtain CD45- cells, and used both 10X and Smart-seq2 to perform scRNA-seq analysis. Following the standard experimental protocols, we obtained 10X data for 1,338, 1,305, 746, and 5,282 cells for LT, MT, NT, and PT tissues, respectively, and obtained Smart-seq2 data for 94, 183, 189, and 135 cells for the corresponding tissues (Table S1). Bulk RNA-seq data of those four samples were also generated.

We first examined the read counts for each cell derived from both platforms. The average total reads of each cell from Smart-seq2 were 6.2M, 1.7M, 6.3M, and 1.7M for LT, MT, NT, and PT, respectively, whereas 10X obtained relatively lower reads as followings: 59K, 34K, 92K, and 20K for the corresponding tissues respectively (Figure 1A and Figure S1A). For transcriptome analysis, we followed conventional practice and selected uniquely mapped reads in the genome for downstream analysis. The number of uniquely mapped reads was nearly 10-fold higher in Smart-seq2 (Figure S2A). Although, the 3’ ends of genes have been reported to have higher homology than other parts of the genome, leading to increased level of multi-alignments [19], our results showed that the unique mapping ratios were similar, at approximately 80% for both datasets (Figure S2A).

**Figure 1.**
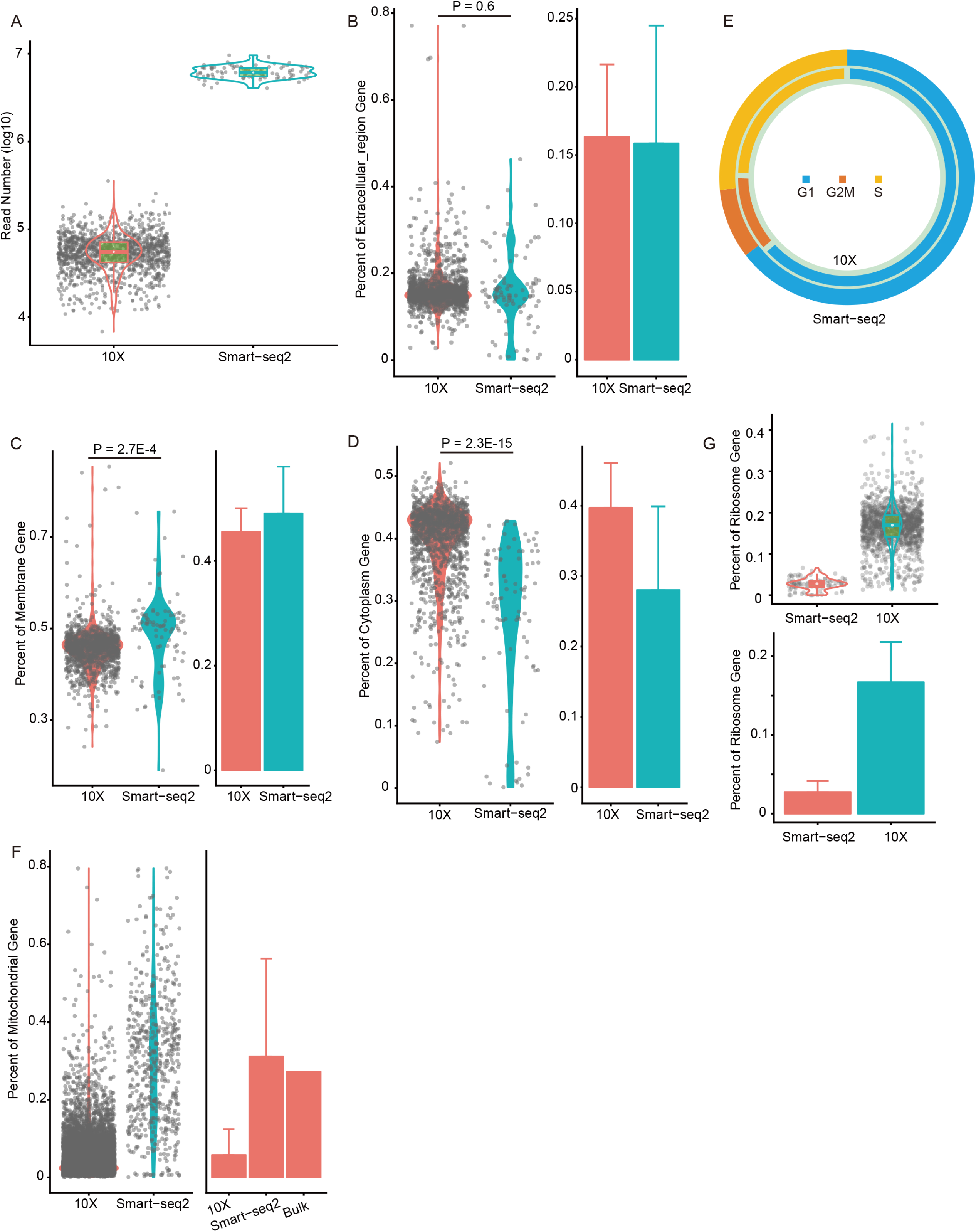
Cell evaluation. **A**. The total reads number of each cell. The proportion of reads of genes in the GO:0005576 “extracellular region” term (**B**), GO:0016020 “membrane” term (**C**), and GO:0005737 “cytoplasm” term (**D**). **E**. The ratio of cells in the G1, G2M, and S phases. The proportion of reads of mitochondrial gene (**F**) and genes in GO:0005840 “ribosome” term (**G**).

As has been reported [20], damaged cells exhibited higher representation of genes in the “membrane” ontology category, but lower representation in the “extracellular region” and “cytoplasm” categories, when compared to high-quality cells. However, we did not observe obvious differences in term of “extracellular region” category between those two scRNA-seq platforms (Figure 1B and Figure S1B). For Smart-seq2, the “membrane” category was over-represented (Figure 1C and Figure S1C) (all P < 1.0E-4, two-sided t-test) and “cytoplasm” category under-represented (Figure 1D and Figure S1D) (all P < 1.0E-10, two-sided t-test), implying more complete lysis of membranes.

Cell cycle has a major impact on gene expression [21], and is an important confounding factor of cell subpopulation classification [22]. We used an established method [23] to classify cells into cell cycle phases based on gene expression (Figure S2B). The distributions of cells in G1, G2/M, and S phases were similar between the two platforms for all samples we studied (Figure 1E and Figure S1E).

### Higher proportion of mitochondrial genes for Smart-seq2 and ribosome-related genes for 10X

One metric we used to examine cell qualities is the proportion of reads mapped to genes in the mitochondrial genome [24]. High levels of mitochondrial reads are indicative of poor quality, likely resulting from increased apoptosis and/or loss of cytoplasmic RNA from lysed cells [20]. Most cells from 10X contained a much lower abundance of mitochondrial genes ranging from 0-15% of their total RNA. By contrast, the mitochondrial proportion from Smart-seq2 was 3.8-10.1 folds higher, at a level similar with bulk RNA-seq data (Figure 1F and Figure S1F). Such high proportions (an average of approximately 30%) by both Smart-seq2 and bulk RNA-seq were likely caused by more thorough disruption of organelle membranes by the Smart-seq2 and the standard bulk RNA-seq protocols than the relatively weak cell lysis procedure by 10X. Abnormally high proportion (such as > 50%) may reflect poor cell quality from Smart-seq2 in this study. However, caveats should be considered when examining mitochondrial genes, because naturally larger mitochondrial proportions can be expected from certain cells such as cardiomyocytes (58-86%) [25] or those in apoptosis [20].

Ribosome-related genes (genes in “ribosome” GO term) accounted for a large portion of detected transcripts by 10X, 3.6-8.2 folds higher than Smart-seq2 data (Figure 1G and Figure S1G). Indeed, 10X detected genes were enriched in the “ribosome” GO term, rather than ribosomal DNA (rDNA). The proportion of sequencing reads assigned to rDNA were only 0.03-0.4% in 10X, significantly lower than those by Smart-seq2 (10.2-28.0%). Few reads were uniquely mapped among those reads (Figure S1H), therefore removing non-uniquely mapped reads was essential to minimizing rDNA interference in Smart-seq2.

### 10X detected a higher proportion of lncRNA and Smart-seq2 identified more lncRNA as highly variable genes

Despite that both Smart-seq2 and 10X followed the poly-A enrichment strategy, approximately 10-30% of all detected transcripts were from non-coding genes (Figure 2A and Figure S3A), with lncRNA accounting for 2.9-3.8% in Smart-seq2 and relatively higher (6.5-9.6%) in 10X (Figure 2B and Figure S3B). In total, protein-coding genes and lncRNA accounted for 80.5-92.6% of all detected transcripts for Smart-seq2, and 77.4-99.2% for 10X. Other classes of RNAs and/or their precursor were also detected with a great variance among experiments. Among protein-coding genes, the proportions of house-keeping (HK) genes and transcriptional factors (TFs) were 1.7-2.5 and 1.1-1.4 folds higher in 10X, respectively (Figure 2C–2D and Figure S3C-S3D).

**Figure 2.**
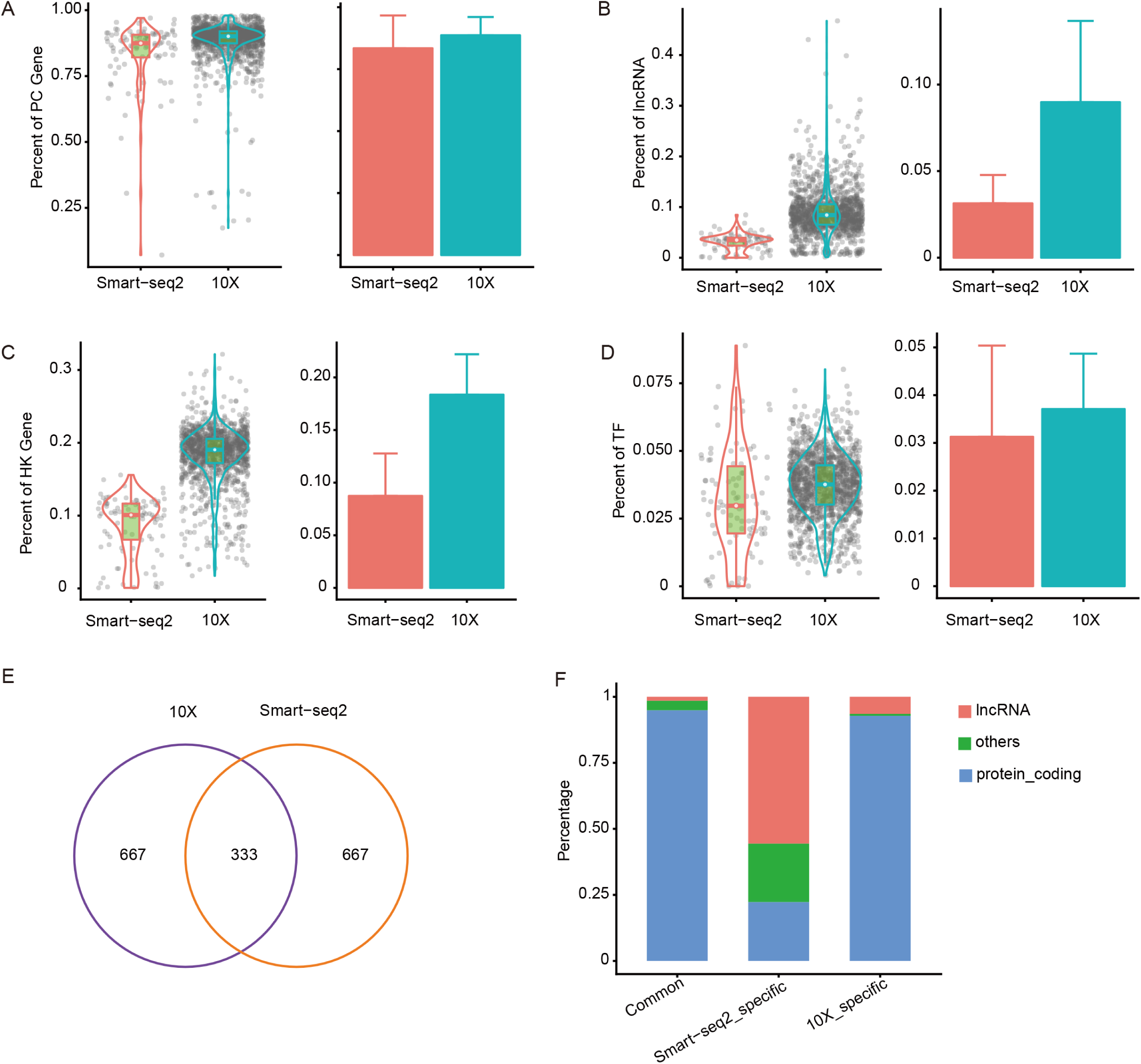
Comparison of lncRNA. The ratio of reads of protein coding (PC) genes (**A**), lncRNA (**B**), house-keeping (HK) genes (**C**), transcription factors (TFs) (**D**). Overlap of highly variable genes (HVGs) identified from 10X and Smart-seq2 (**E**). Types of HVGs (**F**).

One common method to cluster in scRNA-seq datasets was to identify highly variable gene (HVG) [26, 27], which assumed that large variation in gene expression across cells mainly come from biological difference rather than technical noise. We selected the top 1,000 HVGs, and found 333 HVGs shared between two platforms (Figure 2E). Smart-seq2 specific HVGs only enriched two KEGG pathways, while 10X specific HVGs enriched 34 pathways, including common pathways in cancer, such as “PI3K−Akt signaling pathway” (Figure S3E), suggesting that HVGs identified by 10X were more conducive to understanding biological difference among samples. Protein-coding genes accounted for 94.9%, 22.3%, and 92.8% of shared, Smart-seq2 specific, and 10X specific HVGs, respectively (Figure 2F). Huge differences in HVGs come from the lncRNA which has been previously shown to be expressed with biological function in scRNA-seq [19]. The enrichment of lncRNA in Smart-seq2-specific HVGs, which resulted in a few enriched KEGG pathways, may be caused by specific sub-populations which predominantly expressed those lncRNA [28, 29]. The possible reasons may lead to less lncRNA identified as HVGs in 10X as follows: lncRNA was detected at much lower levels than protein-coding genes [30, 31], and higher dropout ratio.

### Smart-seq2 detected more genes and 10X identified more cell clusters

We first assessed the gene-detection sensitivity, represented as the number of detected genes (TPM > 0 or UMI > 0) per cell [32]. Smart-seq2 had significantly higher sensitivity, capturing an average of 5,713, 4,761, 4,079, and 3,860 genes per cell for LT, MT, NT, and PT, respectively, compared to 2,682, 1,853, 2,123, and 1,104 genes for 10X, respectively (Figure 3A and Figure S4A). In total, more than 25,000 genes were covered from each sample by Smart-seq2; however, despite a magnitude more cells captured by 10X, approximately 20% genes were still dropped out (Figure 3B and Figure S4B). For detected genes, Smart-seq2 data showed a unimodal distribution with few low-expressed genes detected in all cells. By contrast, 10X data showed an obvious bimodal distribution due to a large number of genes with near-zero expression (Figure 3C and Figure S4C), suggesting higher noise or random capture of mRNA at very low expression level.

**Figure 3.**
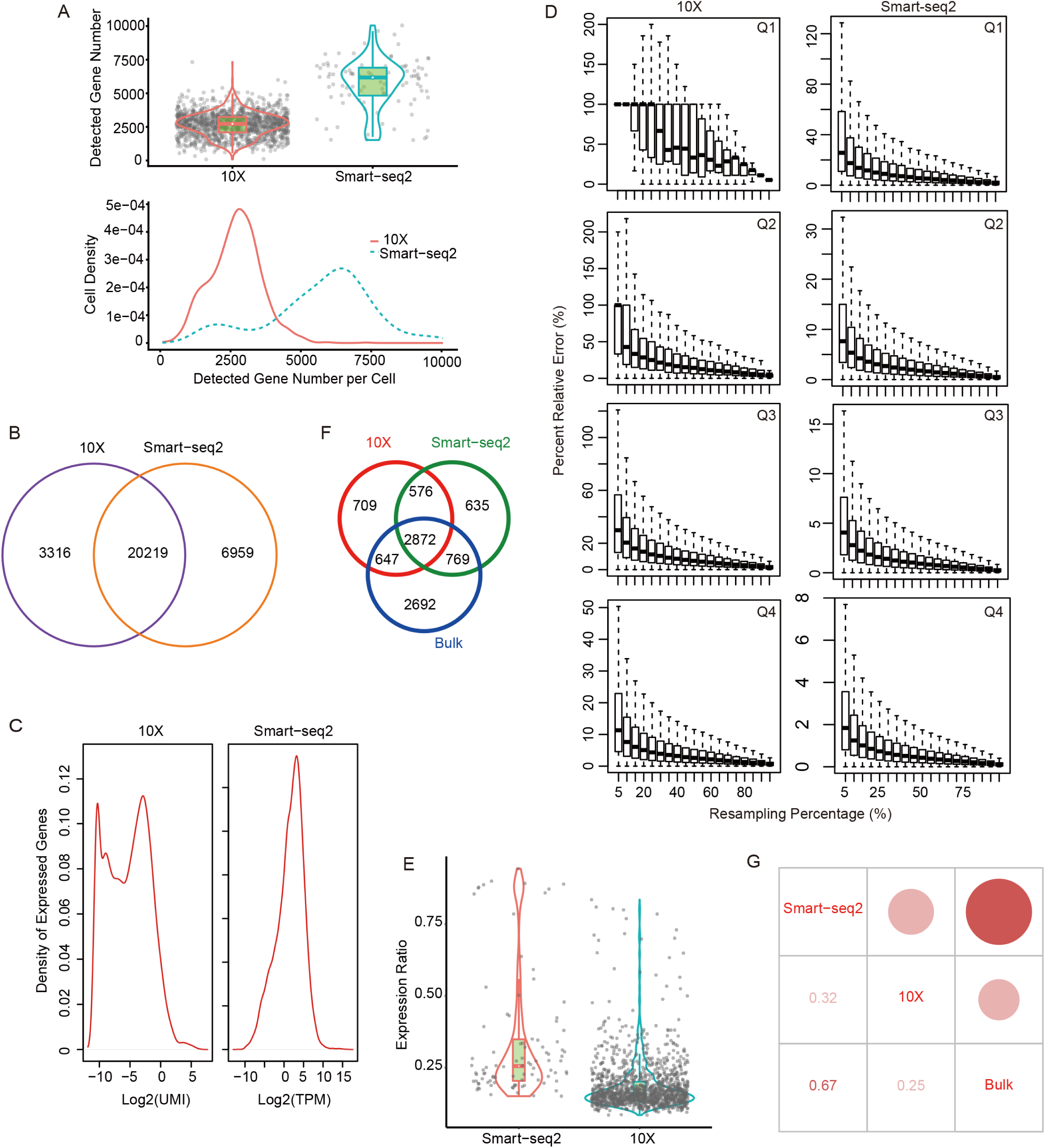
Comparison of detected genes and their expression. **A**. The number of detected genes in every cell. **B**. Overlap of all detected genes between 10X and Smart-seq2. **C**. Distribution of detected genes based on their expression levels. **D**. Saturation analysis by resampling a series of subsets of total reads. **E**. The ratio of reads of the top10 high expressed genes. **F**. Overlap of the top25% high expressed genes among 10X, Smart-seq2, and bulk RNA-seq. **G**. Correlation of expression of common detected genes among 10X, Smart-seq2, and bulk RNA-seq.

To examine the expression dynamic ranges covered by each platform, we determined the expression levels reaching saturation. All genes were divided into four quartiles by expression values. While sequencing depths of all four quartiles were saturated for Smart-seq2, only upper two quartiles were adequate for 10X (Figure 3D and Figure S4D), suggesting that Smart-seq2 has advantages in detecting genes at low expressed levels. Meanwhile, the top 10 most highly expressed genes accounted for 33.0-38.5% of total counts in Smart-seq2 and 18.4-33.0% in 10X (Figure 3E and Figure S4E). Those 10 genes were dominated by mitochondrial genes, especially in Smart-seq2. Moreover, bulk RNA-seq data showed strikingly similar results to Smart-seq2 (Table S2).

We next determined if the two platforms covered different sets of genes. For any given sample, approximately 2/3 of genes present in the upper quartile were shared between the two platforms, leaving the remaining 1/3 genes distinct (Figure 3F and Figure S4F). Analysis of the distinct genes represented indicated that 5.6% of 10X detected genes had full KEGG annotation, whereas only 2.7% of Smart-seq2 detected genes were annotated (Table S3). Thus, Smart-seq2 is better equipped at finding genes with unknown functions. In addition, Smart-seq2 shared more genes with bulk RNA-seq (Figure 3F and Figure S4F). PCC of each gene between bulk RNA-seq and the averaged Smart-seq2 single cell output was higher (Figure 3G and Figure S4G), again showing more similarity between Smart-seq2 and bulk RNA-seq.

HVGs were used to cluster cells into putative subpopulations, which was one of the most common goals of an scRNA-seq experiment. 11 clusters were identified in 10X using Seurat (version 2.3.4) [33]. By applying conventional cell markers, those clusters were annotated as fibroblasts, epithelial cells, endothelial cells, and two special clusters: “hepatocyte” and “malignant cell”, which highly expressed their respective markers, such as, ALB and SERPINA1 in hepatocyte, STMN1, H2AFZ, CKS1B, and TUBA1B in malignant cells [34, 35] (Figure 4A). By contrast, only five clusters were identified in Smart-seq2 due to limited cell number, these clusters were annotated as epithelial cells, endothelial cells and fibroblasts (Figure 4B). Four clusters of tumor fibroblasts were identified in 10X: cluster 0, cluster 2, cluster 5 and cluster 10 (Figure 4A). Cluster 0 cells showed fibroblasts signatures (RGS5 and NDUFA4L2), cluster 2 cells had strong expression of CAF (cancer associated fibroblasts) cell markers (LUM, SFRP4, and COL1A1), cluster 5 cells expressed myofibroblasts markers (MYH11, TAGLN, and ACTA2). We also highlight a fibroblasts cluster (cluster 10) with a striking enrichment for mitochondrial genes (MT-ND2, MT-CO3, and MT-CO2). Smart-seq2 only identified two fibroblasts subtypes, with cluster 2 cells expressing fibroblasts signatures (RGS5 and NDUFA4L2), and cluster 4 cells showing CAF markers (LUM, DCN, and FBLN1).

**Figure 4.**
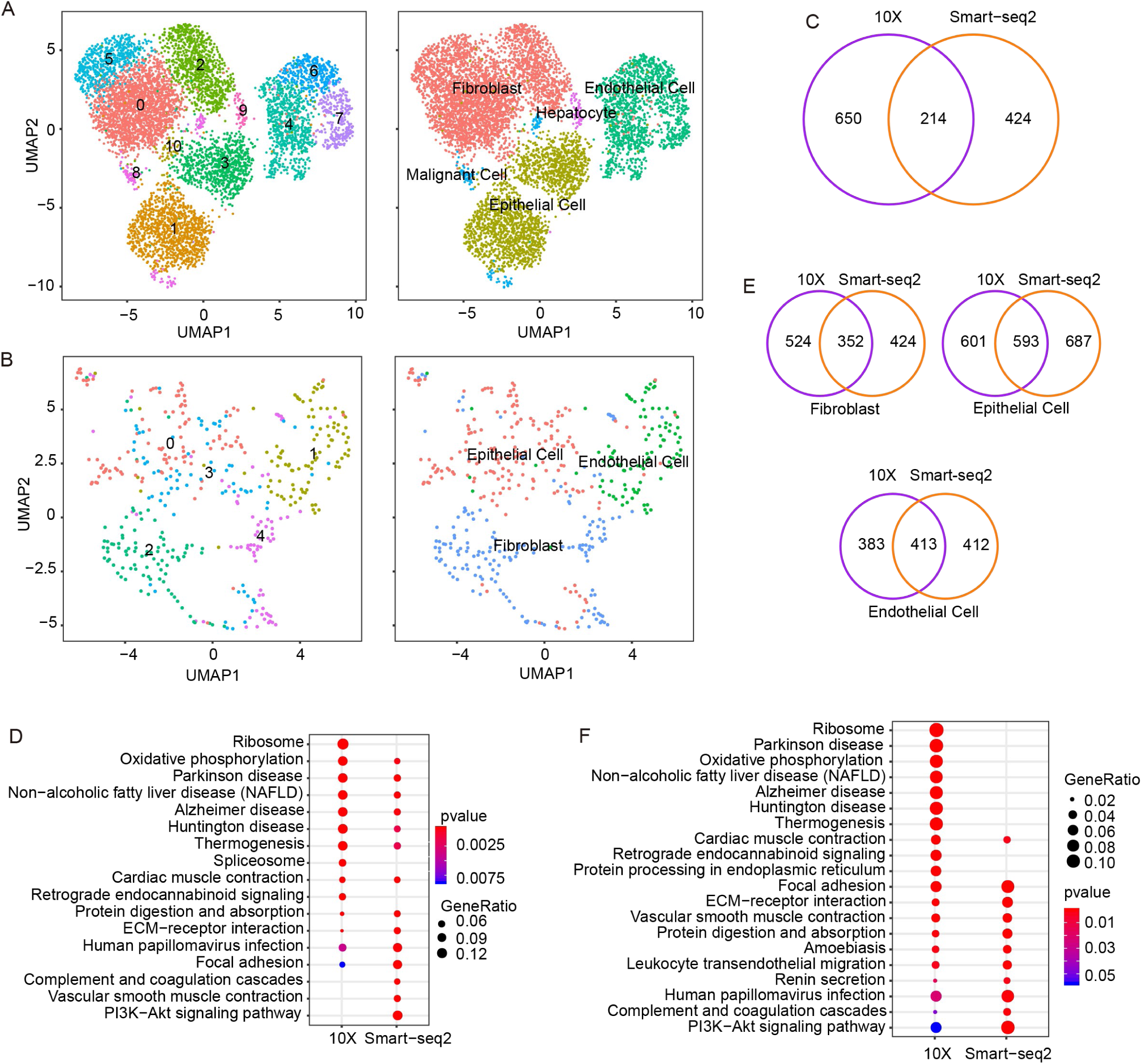
Results of cells clustering and differentially expressed genes (DEGs) Cell clustering results for 10X (**A**) and Smart-seq2 (**B**). **C**. Overlap of DEGs of LT (liver tumor) sample with other three samples identified by 10X and Smart-seq2. Comparison of KEGG enrichment results of LT sample (**D**) and fibroblasts (**F**). **E**. Overlap of DEGs of each cell type compared with remaining types between 10X and Smart-seq2.

We next examined if the two platforms covered different sets of differentially expressed genes (DEGs). We first identified DEGs within each sample compared to all other samples (Figure 4C and Figure S5A). 10X detected more DEGs, and less than 50% of total DEGs were shared between two platforms, leaving the remaining genes distinct. For example, 864 DEGs were identified between LT and other samples using 10X, and 20 KEGG pathways were enriched. Such number were 638 DEGs and 22 pathways for Smart-seq2, respectively. Only 214 DEGs (Figure 4C) and 11 pathways (Figure 4D) were shared. Considering up-regulated DEGs and down-regulated genes separately, less than 50% DEGs were shared between two platforms as well (Figure S5B). Moreover, we observed a few DEGs with conflicting directions (Table S4). We furthermore identified DEGs within each cell type compared to all other cell types (Figure 4E and Figure S5C). The same tendency was also found with several conflicted DEGs (Table S5). Exemplified with fibroblasts, 876 DEGs were identified between fibroblasts and other type cells, and enriched in 30 KEGG pathways in 10X, whereas 776 DEGs identified and 23 pathways enriched in Smart-seq2. Only 352 DEGs (Figure 4E) and 11 pathways (Figure 4F) were shared. In summary, the concordance between DEGs and enriched KEGG pathways by Smart-seq2 and 10X was limited, suggesting that the choice of platform indeed have an impact on the results. Notably, the “Ribosome” pathway was spotted in 10X results (Figure 1G, Figure 4D and 4F, Figure S3E), showing gene detection bias of 10X.

### 10X had higher dropout ratio than Smart-seq2

Dropout events in scRNA-seq can result in many genes undetected and an excess of expression value of zero, leading to challenges in differential expression analysis [21, 36]. The average dropout ratios of majority genes in 10X were 1.3 to 1.4-fold higher for all samples tested (Figure 5A and Figure S6A). For example, the widely used HK gene ACTB had no dropout in Smart-seq2, whereas 2.8-5.9% dropout ratios were observed in 10X. Similarly, GAPDH had dropout ratios from 0-0.67% in Smart-seq2 but 4.2-18.8% in 10X (Figure 5B and Figure S6B).

**Figure 5.**
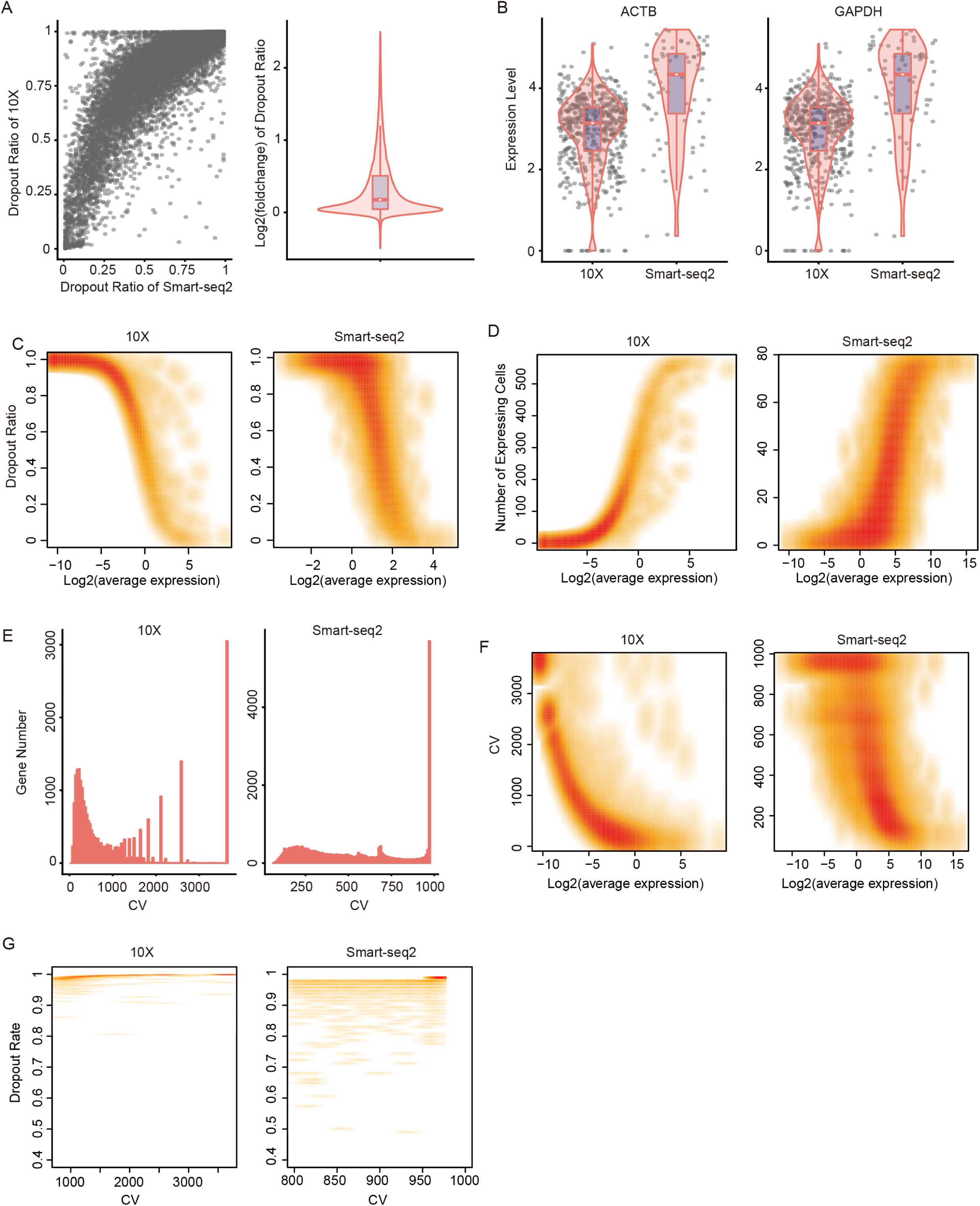
Dropout assessment. **A**. Comparison of dropout ratios between 10X and Smart-seq2. **B**. Two examples of house-keeping genes to show dropout events. **C**. The relationship of dropout ratios and the average expression for each gene. **D**. Number of expressing cells against the average expression of each gene. **E**. CV (coefficient of variation) distribution of each detected gene. **F**. The relationship between CV and gene expression levels. **G**. Dropout ratios of gene with CV more than 800.

The frequency of dropout events was correlated to gene expression levels, which can be fitted by a modified non-linear Michaelis-Menten equation introduced in the M3Drop package (https://github.com/tallulandrews/M3Drop). Genes with lower expression levels had higher dropout ratios (Figure 5C and Figure S6C), consistent with a previous report [37]. Mitochondrial genes were the least likely to be dropped out, especially in Smart-seq2 (Table S6). In both platforms, genes with lower abundance were detected in smaller number of cells, and those genes could lead to higher noise, especially in 10X (Figure 5D and Figure S6D). Because that genes with near-zero expression are noise without enough information for reliable statistical inference [38], removal of them may mitigate noise level and reduce the amount of computation without much loss of information.

We also found that the gene expression coefficient of variation (CV) across cells were associated with dropout ratios. 10X had more genes with large CV than Smart-seq2 (Figure 5E and Figure S6E). While genes with large CV generally had lower expression, especially for 10X (Figure 5F and Figure S6F), genes with larger CV also had higher dropout ratio (Figure S6G). For example, genes with CV larger than 800 had > 80% of dropout ratio in Smart-seq2, near 100% of dropout in 10X (Figure 5G and Figure S6H).

### Difference in capture of gene structural information

We finally evaluated how each of the two platforms capture the gene structural information. We first confirmed that the 10X reads showed a strong bias toward the 3’ ends of mRNAs as expected, while Smart-seq2 reads were more uniformly distributed in the gene bodies (Figure 6A–6B and Figure S7A-S7B). For Smart-seq2, our sequencing depth was adequate for junction detection, evidenced by the number of detected known junctions reaching a plateau (Figure 6C and Figure S7C). The 10X data were not equipped for alternative splicing analysis due to the 3’-bias (Figure 6C and Figure S7C). Nevertheless, 10X still detected non-negligible number of junctions, even though they only accounted for approximately 50% of those junctions detected by Smart-seq2. Although Smart-seq2 data were clearly much more suitable for alternative splicing studies [39, 40], the limited number of splicing junctions detected by 10X might be suitable for certain analyses that rely on junction-based characterization, such as the RNA velocity analysis [41].

**Figure 6.**
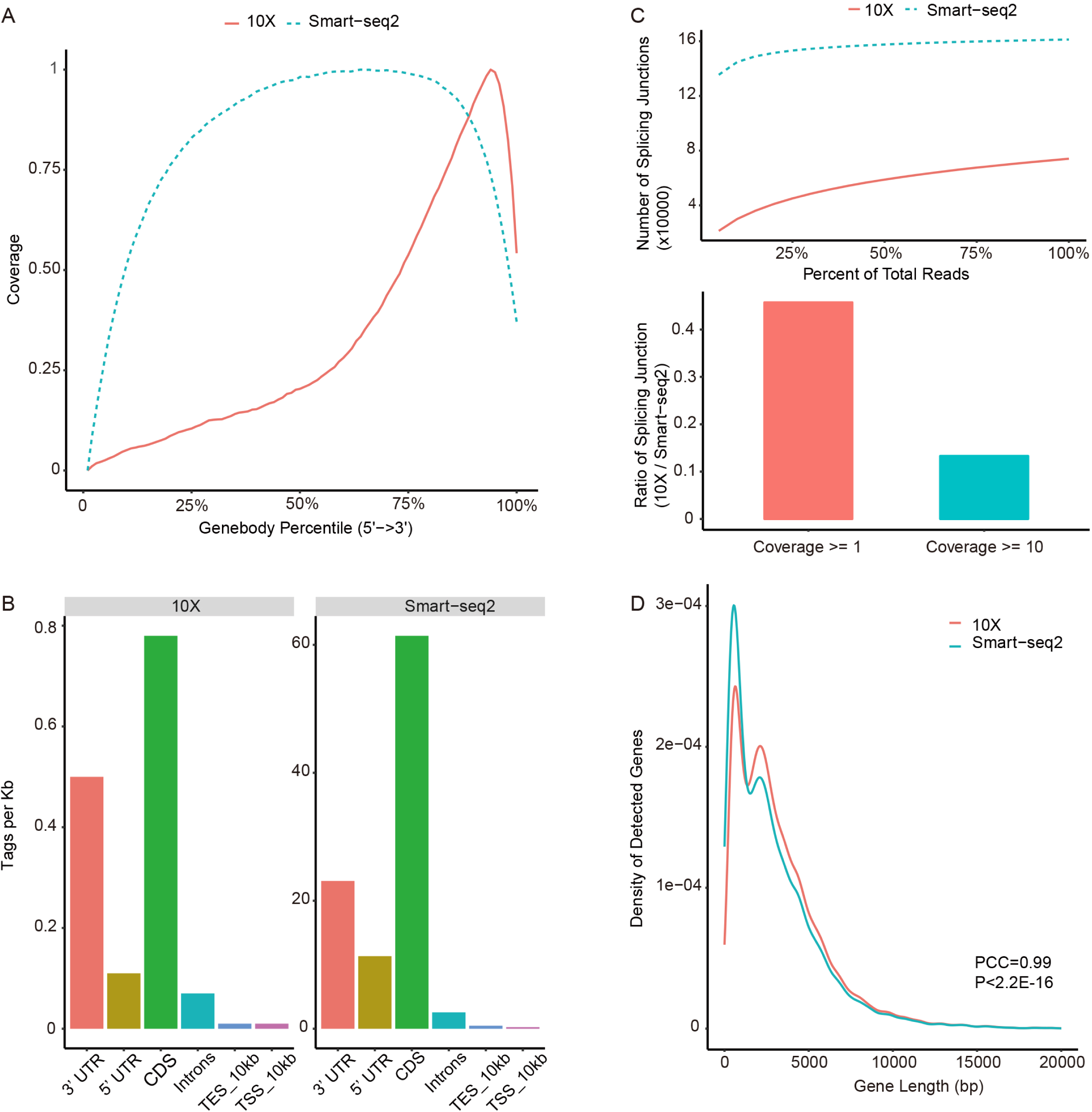
Comparison of gene structural information. **A**. The reads coverage over gene body. **B**. Reads distribution in genome. **C**. Detection of known splice junctions. **D**. Gene length was divided into consecutive 100 bins, we counted the number of detected genes in each bin, PCCs (Pearson correlation coefficients) of gene number between Smart-seq2 and 10X were calculated.

To evaluate whether gene lengths would introduce any bias in either of the platforms, we examined the correlation between the two platforms in terms of gene length and expression level. All calculated PCCs were near perfect for all tested samples (Figure 6D and Figure S7D), demonstrating that mRNA molecular quantification was not influenced by either full-length or 3’ capture strategies.

## Discussion

Here we comprehensively evaluated two scRNA-seq platforms: Smart-seq2 was more sensitive for gene detection, and 10X had more noise and higher dropout ratio. 10X could detect rare cell populations due to high cell throughput. Both platforms had similar results in unique mapping ratio and assigning cells into different cell cycle phase. Smart-seq2 had better performance in detection of genes with low expression levels and of splicing junction. In terms of defining HVGs and detection DEGs, each platform showed unique strength with limited overlap and they could provide complementary information. However, there are some limitations that should be acknowledged in our study. Firstly, the analysis of dropout rates was influenced by the large difference in sequencing depth of those two platforms. Considering an intrinsic property of the two methods, we did not perform downsample to equal sequencing coverage. Secondly, we only sequenced 94-189 cells per sample with the Smart-seq2 protocol, which may reduce the power to detect groups of cells. As has been previously shown, Smart-seq2 libraries should contain ∼70 cells per cluster to achieve decent power [42]. Lastly, UMI counts and read counts have different mean distributions, namely the negative binomial model is a good approximation for UMI counts, and zero-inflated negative binomial for read counts [43], which may impair the CV measure because that CV is linked to the mean gene expression levels.

The advantage of scRNA-seq crucially depends on two parameters: cell number and sample complexity. These two parameters can be designed and chosen based on study objectives. The number of cells is a key determinant for profiling the cell composition. In this study, several hundreds of cells could capture abundant, but not rare, cell types using Smart-seq2. Thousands of cells or more could capture unique cell subtypes in both Smart-seq2 and 10X. Thus, we believe that the range of sample sizes in our comparisons are relevant for other study. In a heterogenous population where the cellular states are transcriptionally distinct and equally distributed, 1,000-2,000 single cells could be sufficient for de novo clustering of the different cell states [44].

However, the cost is still prohibitive for studies that involve hundreds of thousands of cells even at low sequencing depths [7]. It seems a now standard practice to investigate tens of thousands of cells in a published paper. The cost is certainly an important factor for the optimal selection of the cell number. Smart-seq2 is not restricted by cell size, shape, homogeneity, and cell number, and thus is an efficient method to uncover an in-depth characteristic of a rare cell population such as germ cells. However, its overall cost is very high, and the laborious nature and technical variability can be intimidating because the reactions are carried out in individual wells for Smart-seq2 [42]. The huge advantage of 10X is the low cost and high throughput, making it better for complex experiments such as multiple treatments. Although many cells of each sample were added to each channel for 10X in our study, we just obtained 746, 1,305, 1,338, and 5,282 cells by CellRanger (version 2.2, http://www.10xgenomics.com/). 10X cannot guarantee the yield of cells, and cell number may fluctuate wildly among experiments. For example, 60-4,930 cells among 68 samples [45], and 1,052-7,247 cells among 25 samples [46] were obtained in two reports, respectively. The huge variability may come from tissue/cell types, inaccurate estimation of input cell number, or poor conditions and death of cells during experiments. Dataset from a small number of cells is not adequate to reflect fully the biological image [47]. Therefore, the trade-off between Smart-seq2 and 10X should be carefully assessed depending on data throughput and ultimate study objectives.

Samples generally contain a mixture of cells at different phases. However, effects of cell cycle cannot be avoided by simply removing cell cycle marker genes, as the cell cycle can affect many other genes [48, 49]. To date, our results demonstrated that Smart-seq2 and 10X have similar power in assigning cells into different cyclic phases.

The scRNA-seq provides biological resolution that cannot be attained by bulk RNA-seq, at a cost of increased noise [50]. Reliable capture of transcripts into cDNA for sequencing is difficult for the low abundance genes in a single cell, which increases the frequency of dropout events. This was more noticeable in 10X (Figure 5C). Moreover, 10X may capture some ambient transcript molecules that float in droplet due to cell lysis or cell death [19], which also results in noise, however, increased capture single cells could compensate the inefficacy brought by noise and provide a more robust clustering. By contrast, Smart-seq2 had less noise and higher sensitivity but high cost, therefore the sample size attribute in Smart-seq2 and 10X should be established on rigorous design and well-defined rationale.

## Conclusions

Here we comprehensively evaluated two scRNA-seq platforms from the aspects of sensitivity, precision and power: Smart-seq2 was more sensitive for gene detection, and 10X had more noise and higher dropout ratio. 10X could detect rare cell populations due to high cell throughput. Both platforms had similar results in unique mapping ratio and assigning cells into different cell cycle phase. Smart-seq2 had better performance in detection of genes with low expression levels and of splicing junction. In terms of defining HVGs and detection DEGs, each platform showed unique strength with limited overlap and they could provide complementary information.

## Materials and methods

### Sample collection and single cell processing

Tumor tissue of two donors were obtained from about 2cm far from tumor edge, and adjacent normal liver tissues (donor 20170608) were located at least 2cm far from the matched tumor tissue. Those fresh tissue were cut into pieces about 1mm^3^ and digested with MACS tumor dissociation kit for 30min. Suspended cells were filtered with 70_μ_m Cell-Strainer (BD) in the RPMI-1640 medium (Invitrogen), then centrifuged at 400g for 5min, and the supernatant was removed. To lyse red blood cells, pelleted cells were suspended in red blood cell lysis buffer (Solarbio) and incubated on ice for 2min. Finally, cell pellets were resuspended in sorting buffer after washed twice with 1x PBS.

### Single cell RNA-seq

Based on fluorescence activated cell sorting (FACS) analysis (BD Aria III instrument), CD45 (eBioscience, cat. no. 11-0459) was used to separate CD45+ and CD45- cells. Cells were sorted into 1.5mL low binding tubes (Eppendorf) with 50mL sorting buffer, and into wells of 96-well plates (Axygen) with lysis buffer, which contained 1_μ_L 10mM dNTP mix (Fermentas), 1_μ_L 10_μ_M Oligo(dT) primer, 1.9_μ_L 1% Triton X-100 (Sigma) plus 0.1_μ_L 40U/_μ_L RNase Inhibitor (Takara).

For 10X, single cells were processed with the GemCode Single Cell Platform using the GemCode Gel Bead, Chip and Library Kits (10x Genomics, Pleasanton) following the manufacturer’s protocol. Samples were processed using kits pertaining to the V2 barcoding chemistry of 10x Genomics. Estimated 10,000 cells were added to each channel with the average recovery rate 2,000 cells. Libraries were sequenced on Hiseq 4000 (Illumina).

For Smart-seq2, transcripts reverse transcription and amplification were performed according to Smart-seq2’s protocol. We purified the amplified cDNA products with 1x Agencourt XP DNA beads (Beckman), then performed quantification of cDNA of every single cell with qPCR of GAPDH, and fragment analysis using fragment analyzer AATI. To eliminate short fragments (less than 500 bp), cDNA products with high quality were further cleaned using 0.5x Agencourt XP DNA beads (Beckman). The concentration of each sample was quantified using Qubit HsDNA kits (Invitrogen). Libraries were constructed with the TruePrep DNA Library Prep Kit V2 (Vazyme Biotech), and sequenced on Hiseq 4000 (Illumina) in paired-end 150bp.

### Bulk RNA isolation and sequencing

After surgical resection, tissue was firstly stored in RNAlater RNA stabilization reagent (QIAGEN) and kept on ice. Total RNA was extracted using the RNeasy Mini Kit (QIAGEN) according to the manufacturer’s instructions. Concentration of RNA was quantified using the NanoDrop instrument (Thermo), and quality of RNA was evaluated with fragment analyzer (AATI). Libraries were constructed using NEBNext Poly(A) mRNA Magnetic Isolation Module kit (NEB) and NEBNext Ultra RNA Library Prep Kit (NEB), and sequenced on Hiseq 4000 (Illumina) in paired-end 150bp.

### Data reference

We used the GRCH38 human genome assembly as reference, which was downloaded from the Ensembl database (Ensembl 88) (http://asia.ensembl.org). The protein coding genes and lncRNA were categorized based on an Ensembl annotation file in the GTF format. Among those non-coding genes, rRNAs, tRNAs, miRNAs, snoRNAs, snRNA and other known classes of RNAs were excluded, and lncRNA were defined as all non-coding genes longer than 200 nucleotides and not belonging to other RNA categories.

We retrieved the signature genes (extracellular region, cytoplasm, mitochondrion, ribosome, apoptotic process, metabolic process, membrane, and cell cycle) from the gene ontology database (GO:0005576, GO:0005737, GO:0005739, GO:0005840, GO:0006915, GO:0008152, GO:0016020, and GO:0007049, respectively) (http://geneontology.org/). A list of human TFs was obtained from the “Animal Transcription Factor Database” (http://bioinfo.life.hust.edu.cn/AnimalTFDB/).

### Quality control for scRNA

For Smart-seq2, sequenced reads were mapped to GRCH38 using the STAR aligner (version 2.6.0a) with the default parameters. These uniquely mapped reads in the genome were used, and reads aligned to more than one locus were discarded. The expression level of gene was quantified by the TPM value. Genes expressed (TPM > 0) in less than 10 cells were filtered out. Cells were removed according to the following criteria: (1) cells that had fewer than 800 genes and (2) cells that had over 50% reads mapped to mitochondrial genes.

For 10X, an expression matrice of each sample was obtained using the CellRanger toolkit (version 2.2, https://www.10xgenomics.com/). Genes presented (UMI > 0) in less than 10 cells were filtered out. Cells were removed according to the following criteria: (1) cells that had fewer than 500 genes; (2) cells that had fewer than 900 UMI or over 8000 UMI; and (3) cells that had more than 20% of mitochondrial UMI counts.

### CV

The coefficient of variation (CV) is a standardized measure of dispersion of a probability distribution or frequency distribution. It is defined as the ratio of the standard deviation (SD) to the mean, namely CV = 100*SD/mean

### Cell cycle

We used the reported method [23] to classify cells into cell cycle phases based on gene expression. Cells were classified as being in G1 phase if the G1 score is above 0.5 and greater than the G2/M score; in G2/M phase if the G2/M score is above 0.5 and greater than the G1 score; and in S phase if neither score is above 0.5 [51].

### Reads distribution in genome and junction detection

To demonstrate the bias of reads distribution in genome, we calculated reads distribution over genome features, including coding sequence (CDS), 5’-untranslated region (UTR), 3’-UTR, intron, TSS_up_10kb (10kb upstream of transcription start site), and TES_down_10kb (10kb downstream of transcription end site). When genome features were overlapped, they were prioritized as follows: CDS > UTR > Intron > others.

We assessed sequencing depth for splicing junction detection by randomly resampling total alignments with an interval of 5%, and then detected known splice junctions from the reference gene model in GTF format.

### Saturation analysis

We resampled a series of alignment subsets (5%, 10% - 100%) and then calculated RPKM value to assess sequencing saturation, which had been described [52]. “Percent Relative Error” was used to measure how the RPKM estimated from subset of reads (RPKM_est_) deviates from real expression level (RPKM_real_). The RPKM estimated from total reads was used as approximate RPKM_real_: Percent Relative Error = 100 * (| RPKM_est_ – RPKM_real_ |) / RPKM_real_.

### Cell clustering

After filtration, a merged expression matrice of four samples was used for cell clustering by the Seurat package (version 2.3), adapting the typical pipeline [33]. In brief, gene expression was normalized by the “NormalizeData” function. Highly variable genes were calculated with the Find Variable Genes method with the default parameters. Data was scaled with mitochondrial count ratio of a cell for Smart-seq2, with total UMI number and mitochondrial count ratio of a cell for 10X. Those HVGs were used for Canonical Correlation Analysis (CCA), which was used to remove batch effects of patients. Cells were clustered by the “FindClusters” method using the first 20 CCs, and UMAP was used to visualization. Subsequently, cell clusters were annotated manually, based on known markers. Hepatocyte marker genes were ALB and SERPINA1, malignant cell marker genes were STMN1, H2AFZ, CKS1B, and TUBA1B, fibroblast marker genes were RGS5 and NDUFA4L2, CAF (cancer associated fibroblast) marker genes were LUM, SFRP4, DCN, FBLN1 and COL1A1, and myofibroblast marker genes were MYH11, TAGLN, and ACTA2.

### Data visualization and statistics

Microsoft R Open (version 3.5.1, https://mran.microsoft.com/) was used, and ggplot2 package (version 3.1.0) were used to generate data graphs. Data were presented as the mean ± SD in figures. Results of LT (liver tumor) sample were shown in Figures, and corresponding results of other three samples were shown in supplementary files. KEGG pathway enrichment (P < 0.01) were performed using clusterProfiler package (version 3.9.2) [53]. Differentially expressed genes were identified by the “FindMarkers” function (“logfc.threshold” = 0.25 and “min.pct” = 0.25) using the MAST method [54], and P value was adjusted using *bonferroni* correction based on the total number of gene in the dataset, with the thresholds of adjusted P < 0.01.

## Authors’ contribution

ZMZ supervised research. XLW and YH analyzed data. XLW and QMZ drafted the manuscript. QMZ did experiments. ZMZ and XWR revised the manuscript. All authors read and approved the final manuscript.

## Acknowledgements

This work was supported by the National Natural Science Foundation of China (Grant No. 31530036, 81573022, and 31601063).

## Data availability

Data will be accessible publicly at the time of publication.

## Ethics approval

This study was approved by the Ethics Committee of Beijing Shijitan Hospital, Capital Medical University. All patients in this study provided written informed consent for sample collection and data analysis.

## Supplementary material

**Table S1 Cell number of each sample**

**Table S2 List of the most highly expressed genes (Top10)**

**Table S3 KEGG enrichment results of 10X-specific, bulk-specific, and Smart-seq2-specific genes in the top25% list**

**Table S4 DEGs among samples with the change trends conflicting**

**Table S5 DEGs among cell types with the change trends conflicting**

**Table S6 List of genes with zero dropout ratio in a sample**

**Figure S1 Cell evaluation of other three samples**

**A**. The total reads of each cell. The proportion of reads of genes in the GO:0005576 “extracellular region” term (**B**), GO:0016020 “membrane” term (**C**), and GO:0005737 “cytoplasm” term (**D**). **E**. The ratios of cells in the G1, G2M, and S phases. The proportion of reads of mitochondrial gene (**F**) and genes in GO:0005840 “ribosome” term (**G**). **H**. Reads proportion of rDNA.

**Figure S2 Assessment of each cell**

**A**. The unique mapping reads of each sample. **B**. Cell cycle phase scores of each cell.

**Figure S3 Comparison of certain classes of genes**

The expression proportion of protein coding (PC) genes (**A**), lncRNA (**B**), house-keeping (HK) genes (**C**), transcription factors (TFs) (**D**). **E**. KEGG enrichment results of 10X-specific, Smart-seq2-specific, and shared highly variable genes (HVGs).

**Figure S4 Comparison of expression profiles**

**A**. The number of detected genes in every cell. **B**. Overlap of all the detected genes between two platforms. **C**. Distribution of detected genes based on their expression levels. **D**. Saturation analysis. Y axis is “Percent Relative Error” which is used to measures how the RPKM estimated from subset of reads deviates from real expression level. **E**. Percentage of total counts assigned to the top 10 most highly-abundant genes. **F**. Overlap of the top25% high expressed genes among 10X, Smart-seq2, and bulk RNA-seq. **G**. Correlation of common detected genes expression among 10X, Smart-seq2, and bulk RNA-seq.

**Figure S5 Results of differentially expressed genes (DEGs)**

**A**. Overlap of DEGs of remaining samples between Smart-seq2 and 10X. Overlap of Up-regulated and down-regulated DEGs for each sample (**B**) and each cell type (**C**) between Smart-seq2 and 10X.

**Figure S6 Dropout events assessment of other three samples**

**A**. Comparison of dropout ratios between 10X and Smart-seq2. **B**. Two examples of house-keeping genes. **C**. The relationships of dropout ratios and the average gene expression levels. **D**. Number of expressing cells against the average expression for each gene. **E**. CV (coefficient of variation) distribution of each detected gene. **F**. The relationship between CV and gene expression levels. Dropout ratios of genes with CV less than 800 (**G**) and genes with CV more than 800 (**H**).

**Figure S7 Comparison of 3’-end VS full-length capture**

**A**. Reads coverage over gene body. **B**. Reads distribution in genome. **C**. Detection of known splice junctions. **D**. PCC (Pearson correlation coefficient) of gene number in consecutive 100 bins divided by gene lengths between Smart-seq2 and 10X.

## Reference

1. Tang F, Barbacioru C, Wang Y, Nordman E, Lee C, Xu N, Wang X, Bodeau J, Tuch BB, Siddiqui A, et al: mRNA-Seq whole-transcriptome analysis of a single cell. Nat Methods 2009, 6:377–382.

2. Pollen AA, Nowakowski TJ, Shuga J, Wang X, Leyrat AA, Lui JH, Li N, Szpankowski L, Fowler B, Chen P, et al: Low-coverage single-cell mRNA sequencing reveals cellular heterogeneity and activated signaling pathways in developing cerebral cortex. Nat Biotechnol 2014, 32:1053–1058.

3. Zhang L, Yu X, Zheng L, Zhang Y, Li Y, Fang Q, Gao R, Kang B, Zhang Q, Huang JY, et al: Lineage tracking reveals dynamic relationships of T cells in colorectal cancer. Nature 2018, 564:268–272.

4. Halpern KB, Shenhav R, Matcovitch-Natan O, Toth B, Lemze D, Golan M, Massasa EE, Baydatch S, Landen S, Moor AE, et al: Single-cell spatial reconstruction reveals global division of labour in the mammalian liver. Nature 2017, 542:352–356.

5. Grover A, Sanjuan-Pla A, Thongjuea S, Carrelha J, Giustacchini A, Gambardella A, Macaulay I, Mancini E, Luis TC, Mead A, et al: Single-cell RNA sequencing reveals molecular and functional platelet bias of aged haematopoietic stem cells. Nat Commun 2016, 7:11075.

6. Benitez JA, Cheng S, Deng Q: Revealing allele-specific gene expression by single-cell transcriptomics. The International Journal of Biochemistry & Cell Biology 2017, 90:155–160.

7. Svensson V, Vento-Tormo R, Teichmann SA: Exponential scaling of single-cell RNA-seq in the past decade. Nat Protoc 2018, 13:599–604.

8. See P, Lum J, Chen J, Ginhoux F: A Single-Cell Sequencing Guide for Immunologists. Frontiers in Immunology 2018, 9:2425.

9. Picelli S, Björklund ÅK, Faridani OR, Sagasser S, Winberg G, Sandberg R: Smart-seq2 for sensitive full-length transcriptome profiling in single cells. Nature Methods 2013, 10:1096–1098.

10. Picelli S, Faridani OR, Bjorklund AK, Winberg G, Sagasser S, Sandberg R: Full-length RNA-seq from single cells using Smart-seq2. Nature Protocols 2014, 9:171–181.

11. Grun D, van Oudenaarden A: Design and Analysis of Single-Cell Sequencing Experiments. Cell 2015, 163:799–810.

12. Stegle O, Teichmann SA, Marioni JC: Computational and analytical challenges in single-cell transcriptomics. Nat Rev Genet 2015, 16:133–145.

13. Shalek AK, Satija R, Shuga J, Trombetta JJ, Gennert D, Lu D, Chen P, Gertner RS, Gaublomme JT, Yosef N, et al: Single-cell RNA-seq reveals dynamic paracrine control of cellular variation. Nature 2014, 510:363–369.

14. Soneson C, Robinson MD: Bias, robustness and scalability in single-cell differential expression analysis. Nat Methods 2018, 15:255–261.

15. Islam S, Zeisel A, Joost S, La Manno G, Zajac P, Kasper M, Lonnerberg P, Linnarsson S: Quantitative single-cell RNA-seq with unique molecular identifiers. Nat Methods 2014, 11:163–166.

16. Fan J, Salathia N, Liu R, Kaeser GE, Yung YC, Herman JL, Kaper F, Fan JB, Zhang K, Chun J, Kharchenko PV: Characterizing transcriptional heterogeneity through pathway and gene set overdispersion analysis. Nature Methods 2016, 13:241–244.

17. Svensson V, Natarajan KN, Ly LH, Miragaia RJ, Labalette C, Macaulay IC, Cvejic A, Teichmann SA: Power analysis of single-cell RNA-sequencing experiments. Nat Methods 2017, 14:381–387.

18. Baran-Gale J, Chandra T, Kirschner K: Experimental design for single-cell RNA sequencing. Brief Funct Genomics 2018, 17:233–239.

19. Azizi E, Carr AJ, Plitas G, Cornish AE, Konopacki C, Prabhakaran S, Nainys J, Wu K, Kiseliovas V, Setty M, et al: Single-Cell Map of Diverse Immune Phenotypes in the Breast Tumor Microenvironment. Cell 2018, 0.

20. Ilicic T, Kim JK, Kolodziejczyk AA, Bagger FO, McCarthy DJ, Marioni JC, Teichmann SA: Classification of low quality cells from single-cell RNA-seq data. Genome Biol 2016, 17:29.

21. Wagner A, Regev A, Yosef N: Revealing the vectors of cellular identity with single-cell genomics. Nat Biotechnol 2016, 34:1145–1160.

22. Zheng C, Zheng L, Yoo JK, Guo H, Zhang Y, Guo X, Kang B, Hu R, Huang JY, Zhang Q, et al: Landscape of Infiltrating T Cells in Liver Cancer Revealed by Single-Cell Sequencing. Cell 2017, 169:1342–1356 e1316.

23. Scialdone A, Natarajan KN, Saraiva LR, Proserpio V, Teichmann SA, Stegle O, Marioni JC, Buettner F: Computational assignment of cell-cycle stage from single-cell transcriptome data. Methods 2015, 85:54–61.

24. Bacher R, Kendziorski C: Design and computational analysis of single-cell RNA-sequencing experiments. Genome Biol 2016, 17:63.

25. Gladka Monika M, Molenaar B, de Ruiter H, van der Elst S, Tsui H, Versteeg D, Lacraz Grègory PA, Huibers Manon MH, van Oudenaarden A, van Rooij E: Single-Cell Sequencing of the Healthy and Diseased Heart Reveals Cytoskeleton-Associated Protein 4 as a New Modulator of Fibroblasts Activation. Circulation 2018, 138:166–180.

26. Klein AM, Mazutis L, Akartuna I, Tallapragada N, Veres A, Li V, Peshkin L, Weitz DA, Kirschner MW: Droplet barcoding for single-cell transcriptomics applied to embryonic stem cells. Cell 2015, 161:1187–1201.

27. Macosko EZ, Basu A, Satija R, Nemesh J, Shekhar K, Goldman M, Tirosh I, Bialas AR, Kamitaki N, Martersteck EM, et al: Highly Parallel Genome-wide Expression Profiling of Individual Cells Using Nanoliter Droplets. Cell 2015, 161:1202–1214.

28. Liu SJ, Nowakowski TJ, Pollen AA, Lui JH, Horlbeck MA, Attenello FJ, He D, Weissman JS, Kriegstein AR, Diaz AA, Lim DA: Single-cell analysis of long non-coding RNAs in the developing human neocortex. Genome Biol 2016, 17:67.

29. Johnson MB, Wang PP, Atabay KD, Murphy EA, Doan RN, Hecht JL, Walsh CA: Single-cell analysis reveals transcriptional heterogeneity of neural progenitors in human cortex. Nat Neurosci 2015, 18:637–646.

30. Cabili MN, Trapnell C, Goff L, Koziol M, Tazon-Vega B, Regev A, Rinn JL: Integrative annotation of human large intergenic noncoding RNAs reveals global properties and specific subclasses. Genes & Development 2011, 25:1915–1927.

31. Hangauer MJ, Vaughn IW, Mcmanus MT: Pervasive Transcription of the Human Genome Produces Thousands of Previously Unidentified Long Intergenic Noncoding RNAs. Plos Genetics 2013, 9:e1003569.

32. Wu AR, Neff NF, Kalisky T, Dalerba P, Treutlein B, Rothenberg ME, Mburu FM, Mantalas GL, Sim S, Clarke MF, Quake SR: Quantitative assessment of single-cell RNA-sequencing methods. Nature methods 2014, 11:41–46.

33. Satija R, Farrell JA, Gennert D, Schier AF, Regev A: Spatial reconstruction of single-cell gene expression data. Nat Biotechnol 2015, 33:495–502.

34. Lambrechts D, Wauters E, Boeckx B, Aibar S, Nittner D, Burton O, Bassez A, Decaluwe H, Pircher A, Van den Eynde K, et al: Phenotype molding of stromal cells in the lung tumor microenvironment. Nat Med 2018, 24:1277–1289.

35. Puram SV, Tirosh I, Parikh AS, Patel AP, Yizhak K, Gillespie S, Rodman C, Luo CL, Mroz EA, Emerick KS, et al: Single-Cell Transcriptomic Analysis of Primary and Metastatic Tumor Ecosystems in Head and Neck Cancer. Cell 2017, 171:1611–1624 e1624.

36. Hashimshony T, Senderovich N, Avital G, Klochendler A, de Leeuw Y, Anavy L, Gennert D, Li S, Livak KJ, Rozenblatt-Rosen O, et al: CEL-Seq2: sensitive highly-multiplexed single-cell RNA-Seq. Genome Biology 2016, 17:77.

37. Brennecke P, Anders S, Kim JK, Kolodziejczyk AA, Zhang X, Proserpio V, Baying B, Benes V, Teichmann SA, Marioni JC, Heisler MG: Accounting for technical noise in single-cell RNA-seq experiments. Nat Methods 2013, 10:1093–1095.

38. Bourgon R, Gentleman R, Huber W: Independent filtering increases detection power for high-throughput experiments. Proc Natl Acad Sci U S A 2010, 107:9546–9551.

39. Deng Q, Ramskold D, Reinius B, Sandberg R: Single-cell RNA-seq reveals dynamic, random monoallelic gene expression in mammalian cells. Science 2014, 343:193–196.

40. Reinius B, Mold JE, Ramskold D, Deng Q, Johnsson P, Michaelsson J, Frisen J, Sandberg R: Analysis of allelic expression patterns in clonal somatic cells by single-cell RNA-seq. Nat Genet 2016, 48:1430–1435.

41. La Manno G, Soldatov R, Zeisel A, Braun E, Hochgerner H, Petukhov V, Lidschreiber K, Kastriti ME, Lonnerberg P, Furlan A, et al: RNA velocity of single cells. Nature 2018, 560:494–498.

42. Ziegenhain C, Vieth B, Parekh S, Reinius B, Guillaumet-Adkins A, Smets M, Leonhardt H, Heyn H, Hellmann I, Enard W: Comparative Analysis of Single-Cell RNA Sequencing Methods. Molecular Cell 2017, 65:631–643.e634.

43. Chen W, Li Y, Easton J, Finkelstein D, Wu G, Chen X: UMI-count modeling and differential expression analysis for single-cell RNA sequencing. Genome Biol 2018, 19:70.

44. Giladi A, Amit I: Single-Cell Genomics: A Stepping Stone for Future Immunology Discoveries. Cell 2018, 172:14–21.

45. Smillie CS, Biton M, Ordovas-Montanes J, Sullivan KM, Burgin G, Graham DB, Herbst RH, Rogel N, Slyper M, Waldman J, et al: Rewiring of the cellular and inter-cellular landscape of the human colon during ulcerative colitis. bioRxiv 2018:455451.

46. Oetjen KA, Lindblad KE, Goswami M, Gui G, Dagur PK, Lai C, Dillon LW, McCoy JP, Hourigan CS: Human bone marrow assessment by single-cell RNA sequencing, mass cytometry, and flow cytometry. JCI Insight 2018, 3:e124928.

47. Tanay A, Regev A: Scaling single-cell genomics from phenomenology to mechanism. Nature 2017, 541:331–338.

48. Buettner F, Natarajan KN, Casale FP, Proserpio V, Scialdone A, Theis FJ, Teichmann SA, Marioni JC, Stegle O: Computational analysis of cell-to-cell heterogeneity in single-cell RNA-sequencing data reveals hidden subpopulations of cells. Nat Biotechnol 2015, 33:155–160.

49. Skinner SO, Xu H, Nagarkar-Jaiswal S, Freire PR, Zwaka TP, Golding I: Single-cell analysis of transcription kinetics across the cell cycle. Elife 2016, 5:e12175.

50. Marinov GK, Williams BA, McCue K, Schroth GP, Gertz J, Myers RM, Wold BJ: From single-cell to cell-pool transcriptomes: stochasticity in gene expression and RNA splicing. Genome Res 2014, 24:496–510.

51. Lun AT, McCarthy DJ, Marioni JC: A step-by-step workflow for low-level analysis of single-cell RNA-seq data with Bioconductor. F1000Res 2016, 5:2122.

52. Wang L, Wang S, Li W: RSeQC: quality control of RNA-seq experiments. Bioinformatics 2012, 28:2184–2185.

53. Yu GC, Wang LG, Han YY, He QY: clusterProfiler: an R Package for Comparing Biological Themes Among Gene Clusters. Omics-a Journal Of Integrative Biology 2012, 16:284–287.

54. Finak G, McDavid A, Yajima M, Deng J, Gersuk V, Shalek AK, Slichter CK, Miller HW, McElrath MJ, Prlic M, et al: MAST: a flexible statistical framework for assessing transcriptional changes and characterizing heterogeneity in single-cell RNA sequencing data. Genome Biol 2015, 16:278.

