## Supplementary figures and images for "Direct Comparative Analysis of 10X Genomics Chromium and Smart-seq2"

### Supplemental Figure 1

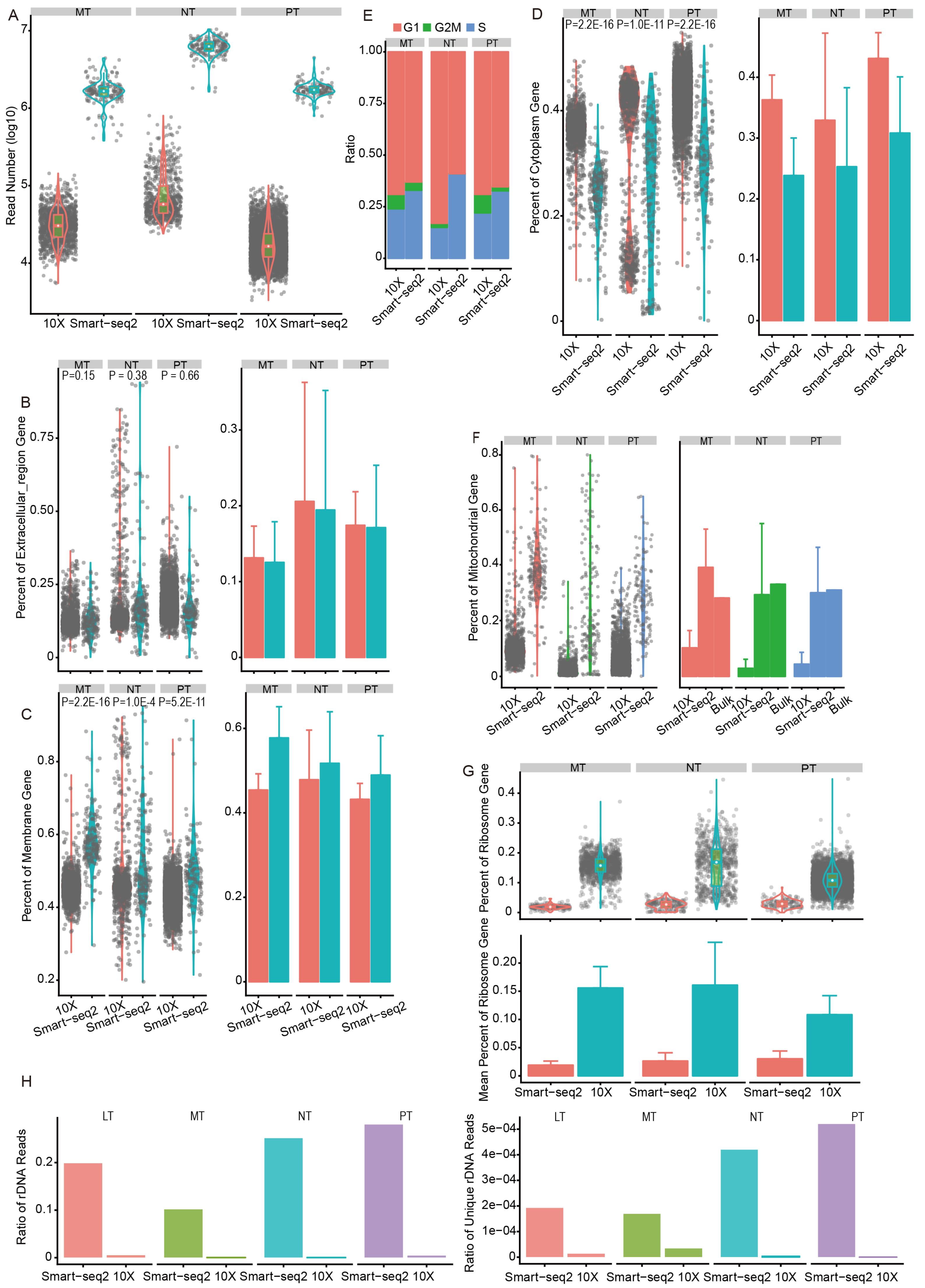

### Supplemental Figure 2

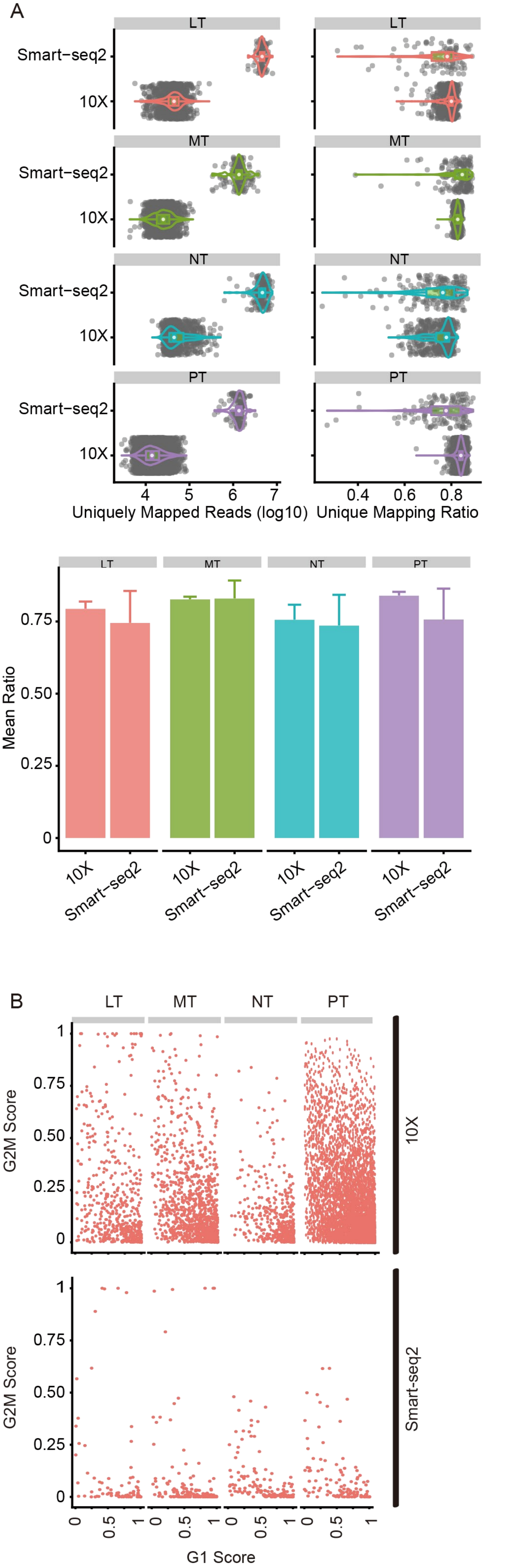

### Supplemental Figure 3

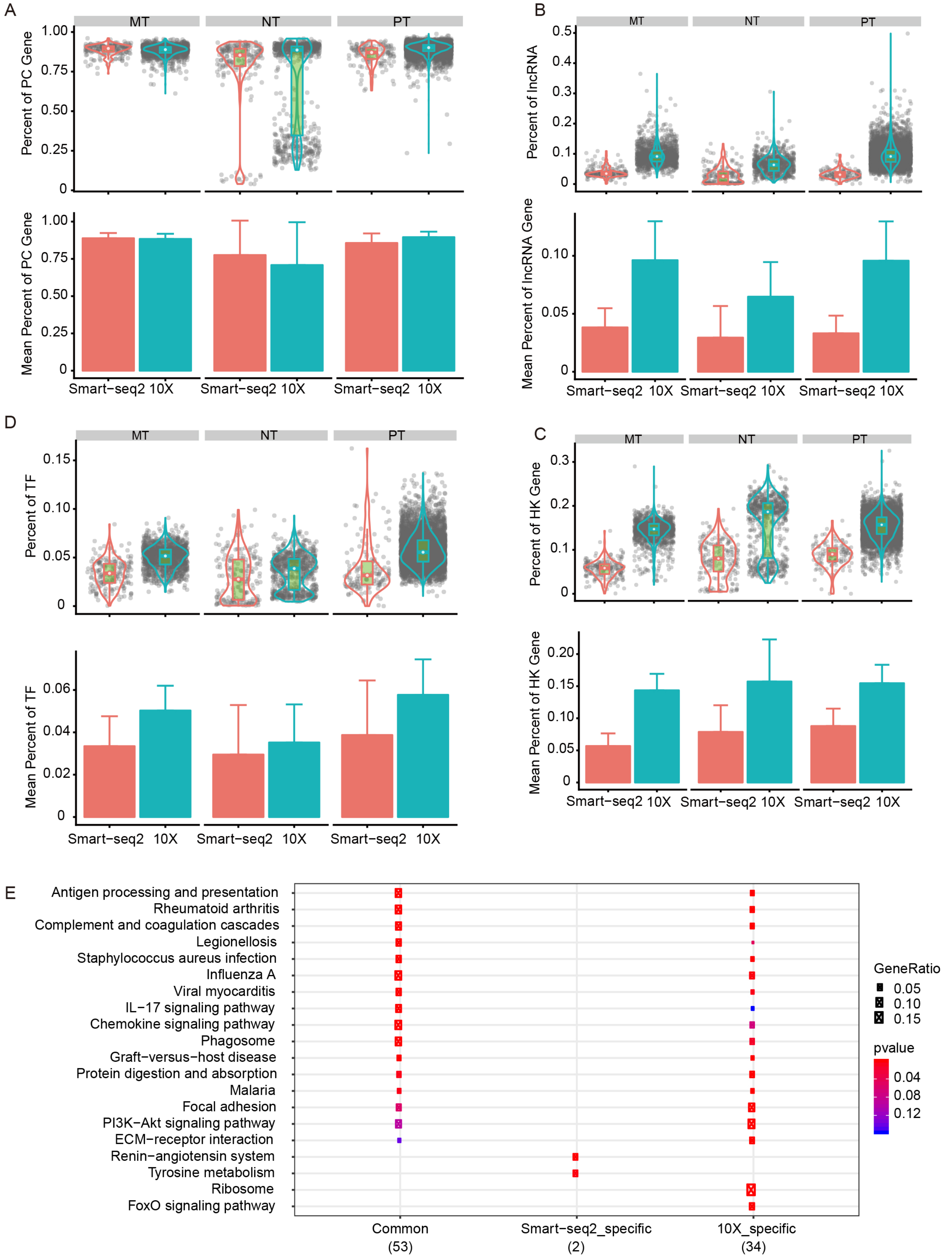

### Supplemental Figure 4

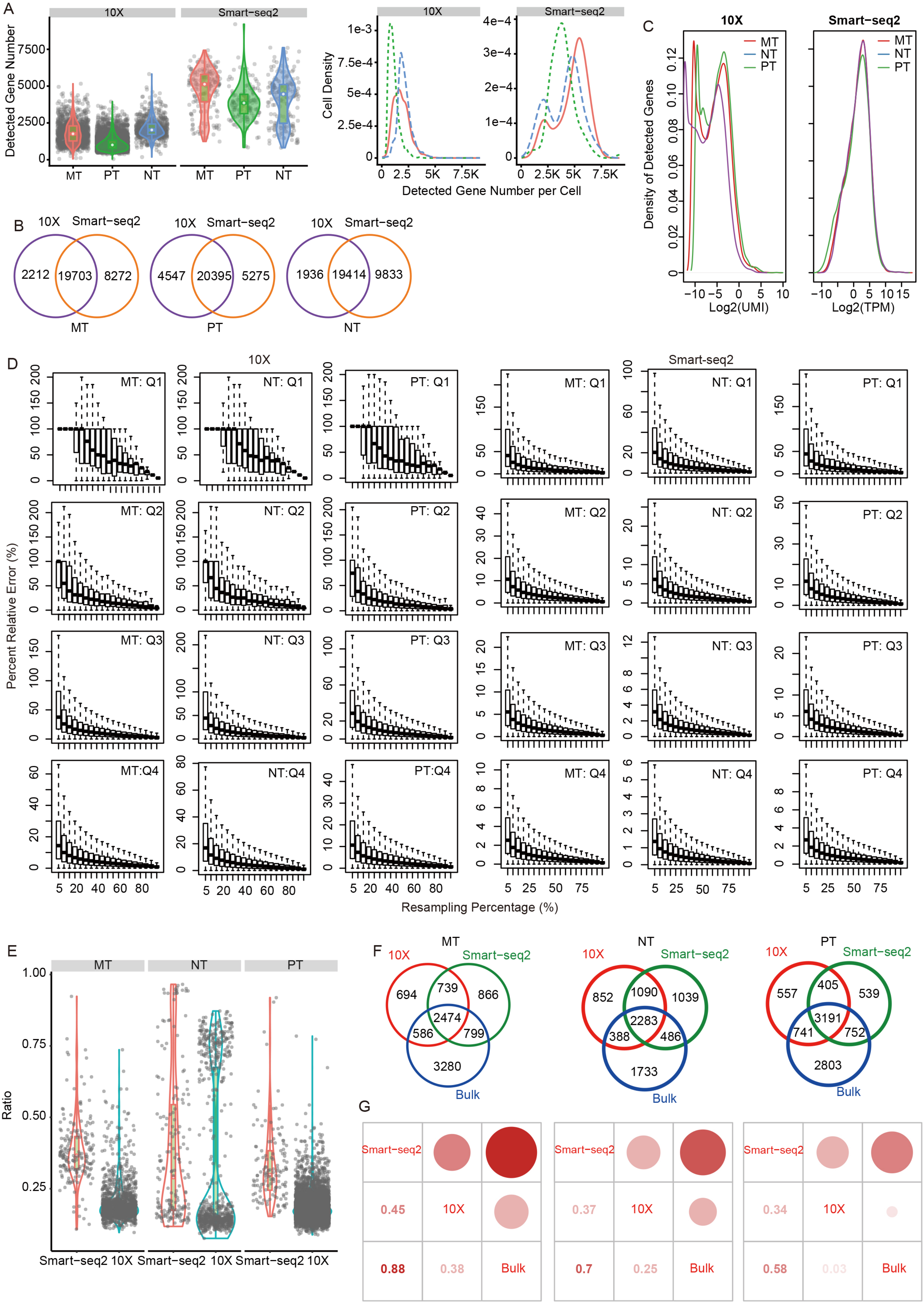

### Supplemental Figure 5

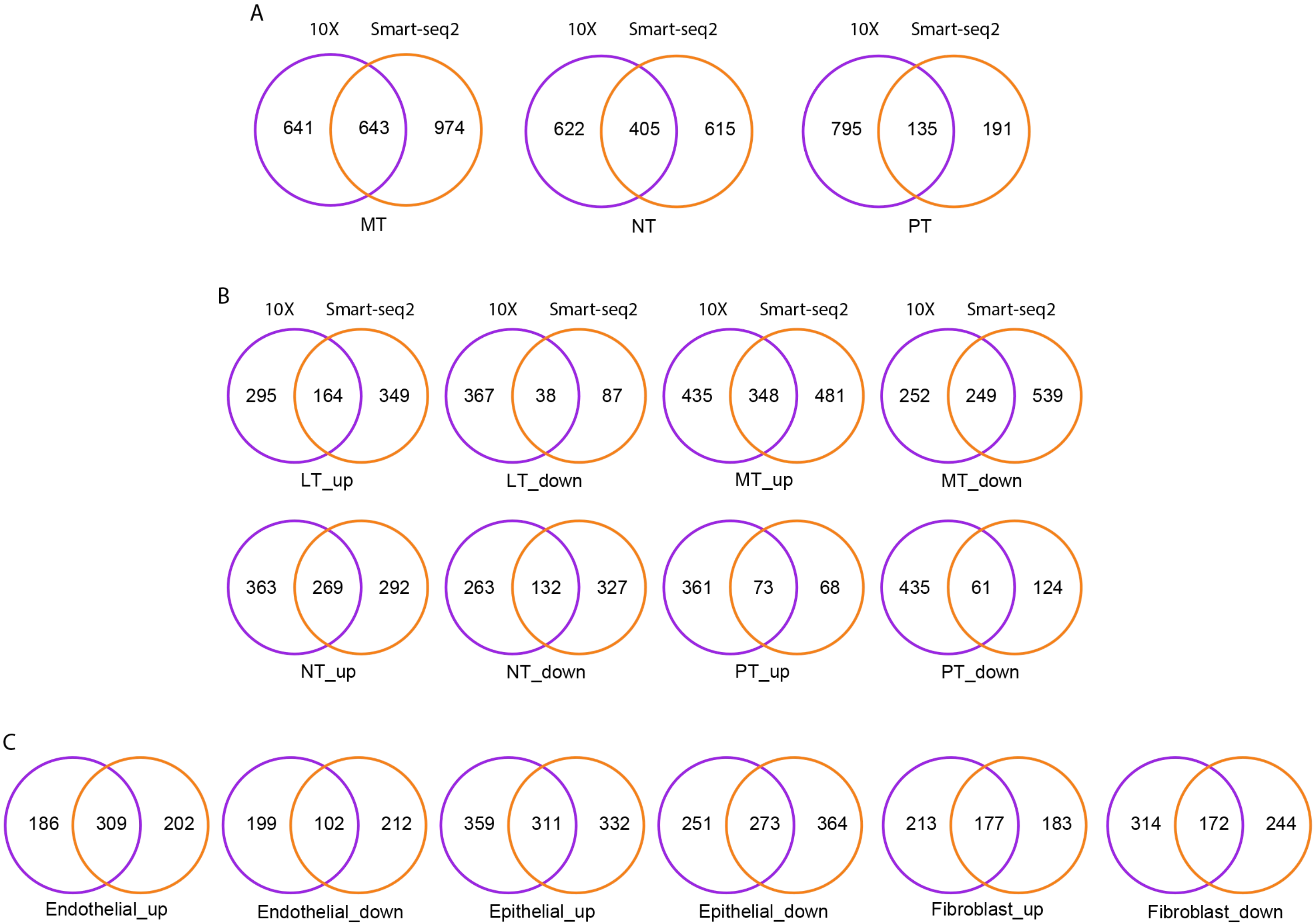

### Supplemental Figure 6

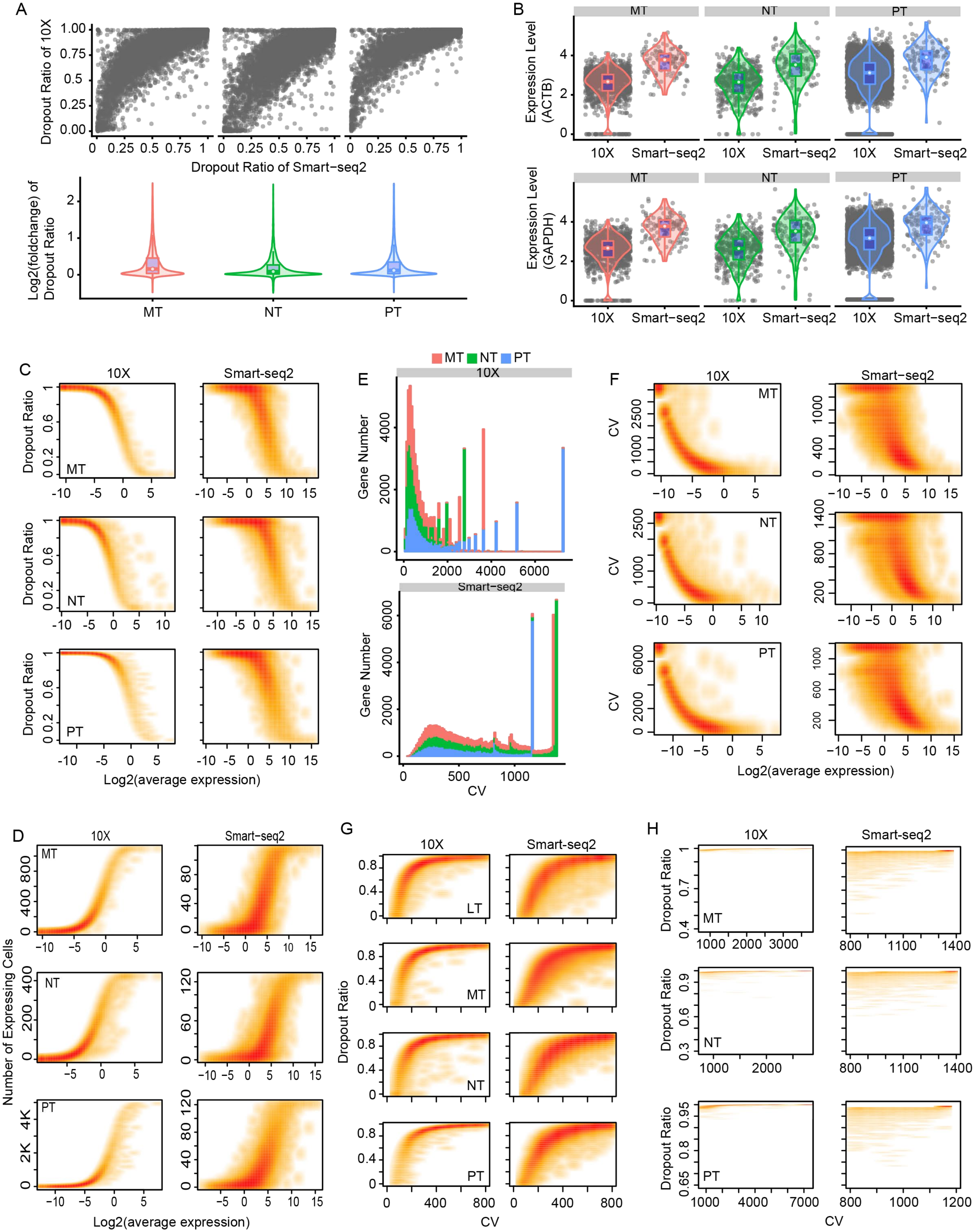

### Supplemental Figure 7

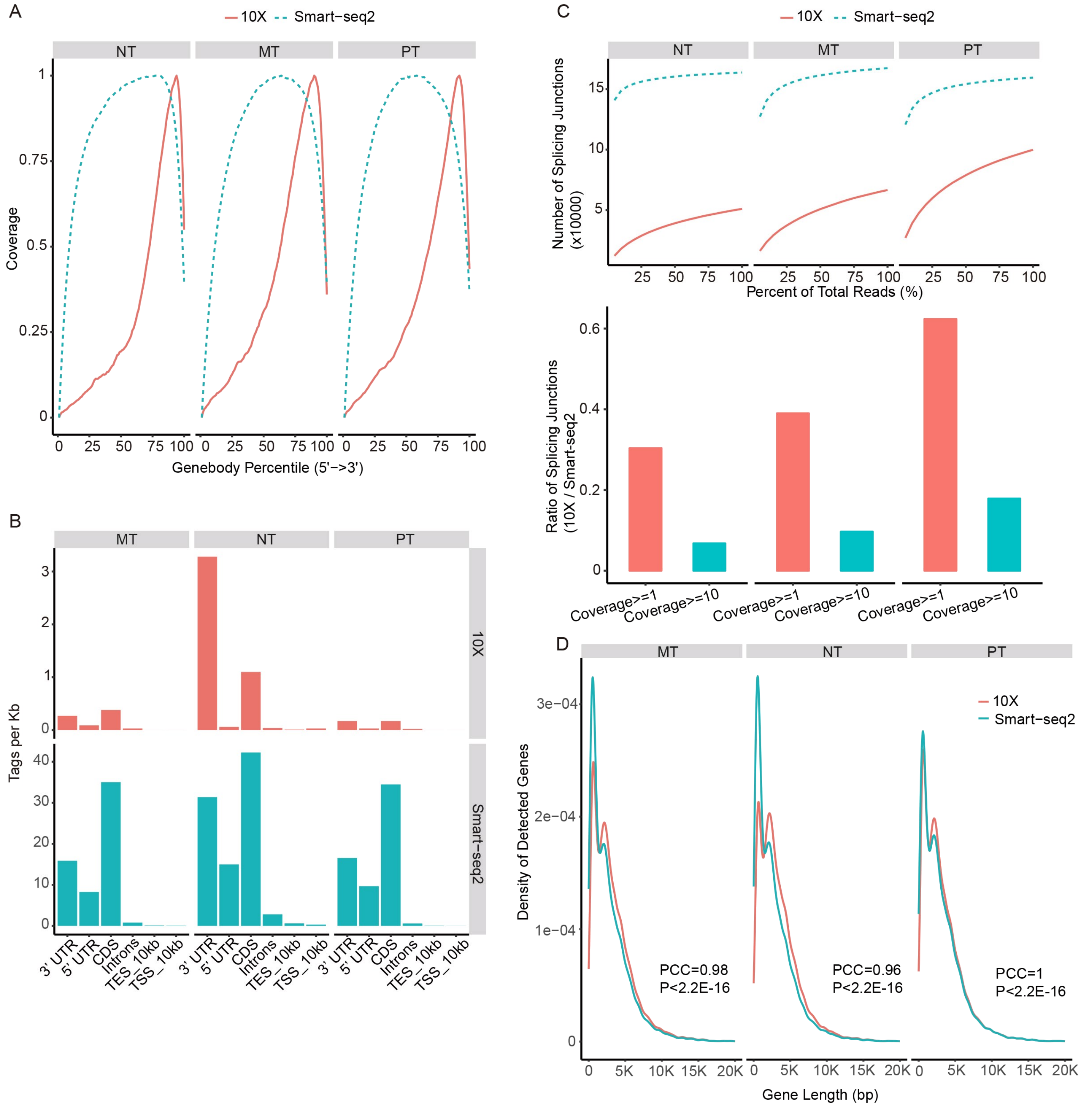
